# HDAC6 inhibition as a mechanism to prevent axon degeneration in the mSOD1^G93A^ mouse model of ALS

**DOI:** 10.1101/2023.08.28.555025

**Authors:** Andrew J Phipps, Samuel Dwyer, Jessica M Collins, Fariha Kabir, Rachel AK Atkinson, Md Anisuzzaman Chowdhury, Lyzette Matthews, Deepika Dixit, Rhiannon S Terry, Jason Smith, Nuri Gueven, William Bennett, Anthony L Cook, Anna E King, Sharn Perry

## Abstract

The loss of upper and lower motor neurons, and their axons is central to the loss of motor function and death in amyotrophic lateral sclerosis (ALS). Due to the diverse range of genetic and environmental factors that contribute to the pathogenesis of ALS, there have been difficulties in developing effective therapies for ALS. One dichotomy emerging in the field is that protection of the neuronal cell soma itself does not prevent axonal vulnerability and degeneration, suggesting the need for targeted therapeutics to prevent axon degeneration. Post-translational modifications of protein acetylation can alter the function, stability and half-life of individual proteins, and can be enzymatically modified by histone acetyltransferases (HATs) and histone deacetyltransferases (HDACs), which add, or remove acetyl groups, respectively. Maintenance of post-translational microtubule acetylation has been suggested as a potential mechanism to stabilise axons and prevent axonal loss and neurodegeneration in ALS. This study has utilized an orally dosed HDAC6 specific inhibitor, ACY-738, prevent deacetylation and stabilize microtubules in the mSOD1^G93A^ mouse model of ALS. Furthermore, co-treatment with riluzole was performed to determine any effects or drug interactions and potentially enhance preclinical research translation. This study shows ACY-738 treatment increased acetylation of microtubules in the spinal cord of mSOD1^G93A^ mice, reduced lower motor neuron degeneration in the lumbar spinal cord of female mice, ameliorated reduction in peripheral nerve axon puncta size, but did not prevent overt motor function decline. The current study also shows peripheral nerve axon puncta size to be partially restored after treatment with riluzole and highlights the importance of co-treatment to measure the potential effects of therapeutics in ALS.

**Highlights:**

- ACY-738 inhibits HDAC6 and leads to increased microtubule acetylation in spinal cord of mSOD1^G93A^ mice.
- ACY-738 treatment reduces lower motor neuron degeneration in the lumbar spinal cord of mSOD1^G93A^ mice.
- ACY-738 treatment restores peripheral nerve axon puncta size of mSOD1^G93A^ mice.
- ACY-738 treatment does not prevent overt motor function decline mSOD1^G93A^ mice.
- Riluzole treatment partially restores peripheral nerve axon puncta size in mSOD1^G93A^ mice.

## Introduction

The neuronal axon is an exquisite structure, forming an integral component of the human nervous system. Despite the axon’s significant role in neuronal communication, it remains understudied in disease and is often overlooked as a target for therapeutic intervention in neurodegenerative diseases such as amyotrophic lateral sclerosis (ALS). Axon degeneration of upper and lower motor neurons is one of the earliest features of the pathophysiology of ALS, leading to muscle loss, sclerosis of the spinal cord, and mortality. Research into peripheral nerves further implicates axon degeneration in the aetiology of ALS, as there is distal to proximal degeneration of axons prior to motor neuron soma loss (Fischer et al., 2004; Fischer & Glass, 2007). In addition, the build-up of proteins such as dynactin, profilin, spastin, neurofilaments, tubulin, and mitochondrial proteins in the axon are precursors to neurodegeneration in ALS (Fischer et al., 2004; Fischer & Glass, 2007; Iwai et al., 2016). Although axon degeneration can occur through independent mechanisms compared to the cell soma, and may even drive neurodegeneration in ALS (Gould et al., 2006), the understanding of mechanisms of axon degeneration, and axonal therapeutic targets in ALS is incomplete.

There are several mechanisms that may contribute to the loss of axons in ALS, including excitotoxicity, axonal accumulation of filamentous protein aggregates, impaired neurofilament structure, and altered post-translational modifications of proteins in the axon (Blizzard et al., 2015; Coleman, 2022; Lefebvre-Omar et al., 2023; Tian et al., 2020; Trist et al., 2022). Given the myriad of suggested drivers of axonal vulnerability in ALS, new therapeutics to prevent axon degeneration would be a valuable novel treatment to prevent neurodegeneration, either alone or in conjunction with known therapies. There have been numerous clinical trials for therapeutics in ALS, however, only two compounds have been approved for treatment by the American Food and Drug Administration; riluzole, and edaverone (Bensimon, Lacomblez, & Meininger, 1994; Cho & Shukla, 2020; Miller, Mitchell, & Moore, 2012). Riluzole’s mechanism of action is to block glutamatergic neurotransmission to reduce ALS associated excitotoxicity (Doble, 1996a), while Edaverone captures peroxyl radicals to reduce nervous system oxidative stress (Rothstein, 2017). These compounds have only shown mild therapeutic benefits, and a 3-6 month extension of life (Bensimon, Lacomblez, & Meininger, 1994; Cho & Shukla, 2020; Miller, Mitchell, & Moore, 2012). The mild neuroprotective nature of these compounds may be attributed to the lack of protection of the axonal structure that also undergoes degeneration through mechanisms potentially independent from the cell body.

Axons have a highly dynamic cytoskeletal structure, composed of actin, intermediate filaments, and microtubules. Microtubules are responsible for axonal transport of cargoes and signals from the cell soma to the synapse, and back. To facilitate axonal transport, microtubules are subject to a wide range of post-translational modifications including phosphorylation, ubiquitination, tyrosination and acetylation, that stabilise or modify the microtubule in response to stimuli or external queues (Kabir et al., 2023). Of particular importance to axonal structure is acetylation, which prevents depolymerisation of tubular subunits, promoting microtubule stabilisation. The process of microtubule acetylation and deacetylation is catalysed through histone acetyltransferase (HATs) and histone deacetyltransferase (HDACs) molecules, respectively. In general, HAT and HDAC molecules are critical in facilitating changes in transcription within the nucleus, however, these molecules also have roles elsewhere in the cell. Sub-classes of HATs and HDACs have non-histone substrates, such as the class IIb HDAC, HDAC6 (Verdel & Khochbin, 1999; Yao & Yang, 2011), which has roles in microtubule acetylation and stabilisation (Osseni et al., 2020). Although HAT enzymes have potential as therapeutics for neurodegenerative diseases, their broad range of substrates within the cell and subsequent lack of substrate specificity, limits their effectiveness. On the other hand, the improved substrate specificity of HDAC enzymes allows for therapeutic potential in neurodegenerative diseases like ALS.

Inhibition of HDACs has been explored as a potential therapeutic in neurodegenerative disease, including Alzheimer’s disease, multiple sclerosis, Huntington’s disease, and ALS (Benoy et al., 2017; Guo et al., 2017; Hanson et al., 2018; Jochems et al., 2014; LoPresti, 2019; Majid et al., 2015; Rossaert et al., 2019). Initial trials of small molecule HDAC inhibitors (HDACi) such as suberoylanilide hydroxamic acid (SAHA) and Trichostatin A have been neuroprotective in models of neurodegenerative disease (Chuang et al., 2009; Saha & Pahan, 2006). In ALS, the pan-HDACi sodium phenylbutyrate prevented dysregulation of apoptotic cell death pathways of the mSOD1^G93A^ mouse model (Ryu et al., 2005), while the pan-HDACi SAHA, prevented oxidative stress-induced motor neuron cell death *in vitro*, and restored the histone deacetylation in SOD1^G86R^ mice (Caroline Rouaux et al., 2007). We have previously shown the pan-HDACi Trichostatin A increases microtubule acetylation in axons and reduces axonal fragmentation after excitotoxic stimuli in murine primary cultures (Hanson et al., 2018). While these studies have shown some beneficial effects from pan-HDACi, improved survival in murine models is poor, potentially due to the inherent limitations of pan-HDACi, such as low blood brain barrier (BBB) permeability, target specificity, and toxicity (Ryu et al., 2005; Sugai et al., 2004; Yoo & Ko, 2011).

More recent studies have focussed on development of novel HDACi candidates with increased specificity and BBB permeability. The targeted inhibition of HDAC6 has been postulated as an effective therapeutic target due to its’ non-histone substrates, and it is actively involved in the post-translational removal of acetylation from microtubules in the axon (reviewed in (Kabir et al., 2023). A promising therapeutic for neurodegenerative disease is a HDAC6 specific inhibitor known as ACY-738 (Jochems et al., 2014). ACY-738 has demonstrated high BBB permeability when administered intraperitoneally or orally, has shown low toxicity in both *in vitro* and *in vivo* studies (Jochems et al., 2014; Majid et al., 2015; Rossaert et al., 2019), and can increase α-tubulin acetylation in the central nervous system *in vivo* (Majid et al., 2015; Rossaert et al., 2019). ACY-738 administration improved axonal transport, reduced hyperphosphorylated tau and improved performance in contextual fear conditioning, and open field performance in the APP/PS1 murine model of Alzheimer’s disease (Majid et al., 2015) and improved short-term memory in an *in vivo* multiple sclerosis model (LoPresti, 2019). Given that ACY-738 has been shown to protect against metabolomic dysfunction (Burg et al., 2021), enhance motor function, prolong lifespan, and restore global histone acetylation in FUS-ALS mice (Rossaert et al., 2019), ACY-738 has therapeutic potential for treating ALS. Furthermore, ACY-738 has been demonstrated to restore axonal transport in FUS-ALS iPSC-derived motor neurons (Guo et al., 2017).

While the effectiveness of ACY-738 as a HDAC6i in FUS-ALS models is promising, it is unknown whether selective HDACi is neuroprotective in other models of ALS. The mSOD1^G93A^ mouse model of ALS is one of the most extensively characterised and used models of ALS pathology, featuring lower motor neuron degeneration, motor function deficits, and axonal degeneration (Gurney et al., 1994). The effectiveness of ACY-738 as a treatment for ALS remains to be tested in the mSOD1^G93A^ model but is essential because of the varied pathological mechanisms driven by different genetic contributors to ALS. Such studies may establish a role for ACY-738 as a drug candidate in those ALS cases with mutant SOD1 variants.

In this study, ACY-738, was used to investigate whether preventing the deacetylation process and stabilizing neuronal microtubules can impede neuronal and axon degeneration and improve motor deficits in the mSOD1^G93A^ mouse model of ALS. Unlike previous research in this area, ACY-738 was simultaneously dosed with riluzole to determine if the two treatments have a combined effect. Given the widespread clinical use of riluzole, it is essential to determine how riluzole interacts with novel therapeutics, given the likelihood of co-treatment. ACY-738 administered to mSOD1^G93A^ mice, acetylated microtubules in the spinal cord, reduced lower motor neuron degeneration and restored peripheral nerve axon puncta size in these animals, whilst riluzole alone also partially restored peripheral nerve axon puncta size. These data have highlighted complex interactions in animals co-treated with riluzole and ACY-738 and have demonstrated some protective effects of HDAC6i on neurodegeneration in mSOD1^G93A^ mice. This research highlights need for further investigations into the role of microtubule acetylation in neurodegeneration.

## Results

### ACY-738 increases microtubule acetylation in the spinal cord of mSOD1^G93A^ mice

To determine that ACY-738 could reach central nervous system targets, a comparison of ACY-738 delivery methods in 12-week-old C57BL/6 mice was performed including intraperitoneal injection of ACY-738 (20mg/kg bodyweight), oral ACY-738 dosing in chow pellets (625mg/kg chow) (Specialty Feeds, WA, Australia), or ACY-738 mixed into a chow porridge daily (625mg/kg porridge). ACY-738 concentrations were chosen based on previous studies utilising the compound to treat neurodegenerative disease in murine models (LoPresti, 2019; Majid et al., 2015; Rossaert et al., 2019). The effectiveness of each treatment was determined by measuring α-tubulin acetylation in the brain and spinal cord. Indirect ELISA assays showed that all the methods of ACY-738 administration significantly increased α-tubulin acetylation in the cortex of 12 week old C57BL/6 mice after acute treatment for 3 days (p<0.05; one-way ANOVA, Tukey’s post-hoc; Figure 1 a), and there was no significant difference in the amount of change of α-tubulin acetylation from differing treatment methods (Figure 1 a). As such, the drug was formulated into pellets for oral dosing of ACY-738 for all future experiments to minimize animal stress and food preparation and was administered to mice for 9 weeks from 12 to 20 weeks of age. The effect of chronic oral dosing of ACY-738 on α-tubulin acetylation was also examined in the spinal cord of mSOD1^G93A^, where ACY-738 administration led to significantly increased acetylated α-tubulin in mSOD1^G93A^ mouse cervical spinal cord, with approximately .75 times the level of acetylated α-tubulin in treated mice, compared to untreated controls (p<0.05; one-way ANOVA, Tukey’s post-hoc; Figure 1b). These data confirmed that ACY-738 effectively increased microtubule acetylation in the spinal cord of mSOD1^G93A^ mice following treatment.

### ACY-738 does not prevent weight loss and motor function decline in mSOD1^G93A^ mice

After establishing delivery methods, drug activity, and capacity to alter HDAC6 activity in neural tissue, mSOD1^G93A^ male and female mice were orally dosed with ACY-738 formulated chow from symptom onset at 12 weeks of age, until late-stage disease at 20 weeks of age (Figure 2-a), which was the chosen ethical endpoint. Weight is an established marker of disease progression in mSOD1^G93A^ mice, due to muscle denervation and resultant atrophy. To investigate the ability of ACY-738 to rescue neuromotor decline associated with ALS, weight was measured weekly from pre-treatment until endpoint.

C57BL/6 mice steadily gained weight from 10 weeks of age until endpoint, whilst male and female mSOD1^G93A^ mice gained weight until 12 weeks of age, at which time weight was maintained or lost until end-point, at 20 weeks of age (Figure 2 a-b). At 20 weeks of age, all groups of mSOD1^G93A^ mice weighed significantly less than untreated C57BL/6 controls (p<0.05; one-way ANOVA, Tukey’s post-hoc; Table 1; Figure 2). Treatment with ACY-738, ACY-738+riluzole, or riluzole alone, did not counteract weight loss in male and female mSOD1^G93A^ mice, relative to wild-type mice, while male and female mice, regardless of treatment, weighed similar to untreated mSOD1^G93A^ controls at 20 weeks of age. There was a main effect of sex at 20 weeks of age, where all male mSOD1^G93A^ mice, regardless of treatment group weighed significantly more than female mice (p<0.05; one-way ANOVA, Tukey’s post-hoc; Table 1, Figure 2). In addition, gastrocnemius muscles were weighed to examine the extent of gross muscle atrophy and denervation at the 20-week end-point. While C57BL/6 mice gastrocnemius muscles weighed significantly more than mSOD1^G93A^ mice at endpoint (p<0.05; one-way ANOVA, Tukey’s post-hoc; Table 1; Figure 2), ACY-738 treatment with, or without riluzole, did not rescue gastrocnemius muscle atrophy in mSOD1^G93A^ mice (Table 1, Figure 2 b-c). Combined, these data suggest ACY-738, and riluzole does not prevent mSOD1^G93A^ body weight loss and muscle denervation.

**Table 1:** Weight of mice at 20 weeks (endpoint) across treatment groups.

| Treatment group | Sex | Sample size | Mean(g) | 95% CI |
| --- | --- | --- | --- | --- |
| mSOD1<sup>G93A</sup> ACY-738 | Male | 13 | 23.44 | 22.14 - 24.15 |
| mSOD1<sup>G93A</sup> ACY-738 + riluzole | Male | 11 | 24.02 | 23.10 - 24.93 |
| mSOD1<sup>G93A</sup> riluzole | Male | 11 | 24.56 | 22.84 - 26.28 |
| mSOD1<sup>G93A</sup> control | Male | 13 | 24.98 | 23.76 - 26.20 |
| C57BL/6 control | Male | 5 | 28.29 | 26.49 - 30.08 |
| mSOD1<sup>G93A</sup> ACY-738 | Female | 18 | 19.36 | 18.67 - 20.05 |
| mSOD1<sup>G93A</sup> ACY-738 + riluzole | Female | 19 | 18.95 | 18.12 - 19.78 |
| mSOD1<sup>G93A</sup> riluzole | Female | 12 | 20.10 | 18.72 - 21.48 |
| mSOD1<sup>G93A</sup> control | Female | 15 | 19.89 | 18.79 - 21.00 |
| C57BL/6 control | Female | 5 | 23.01 | 21.32 - 24.69 |

*Data presented as conditional means and 95% confidence intervals.*

**Figure 1:**
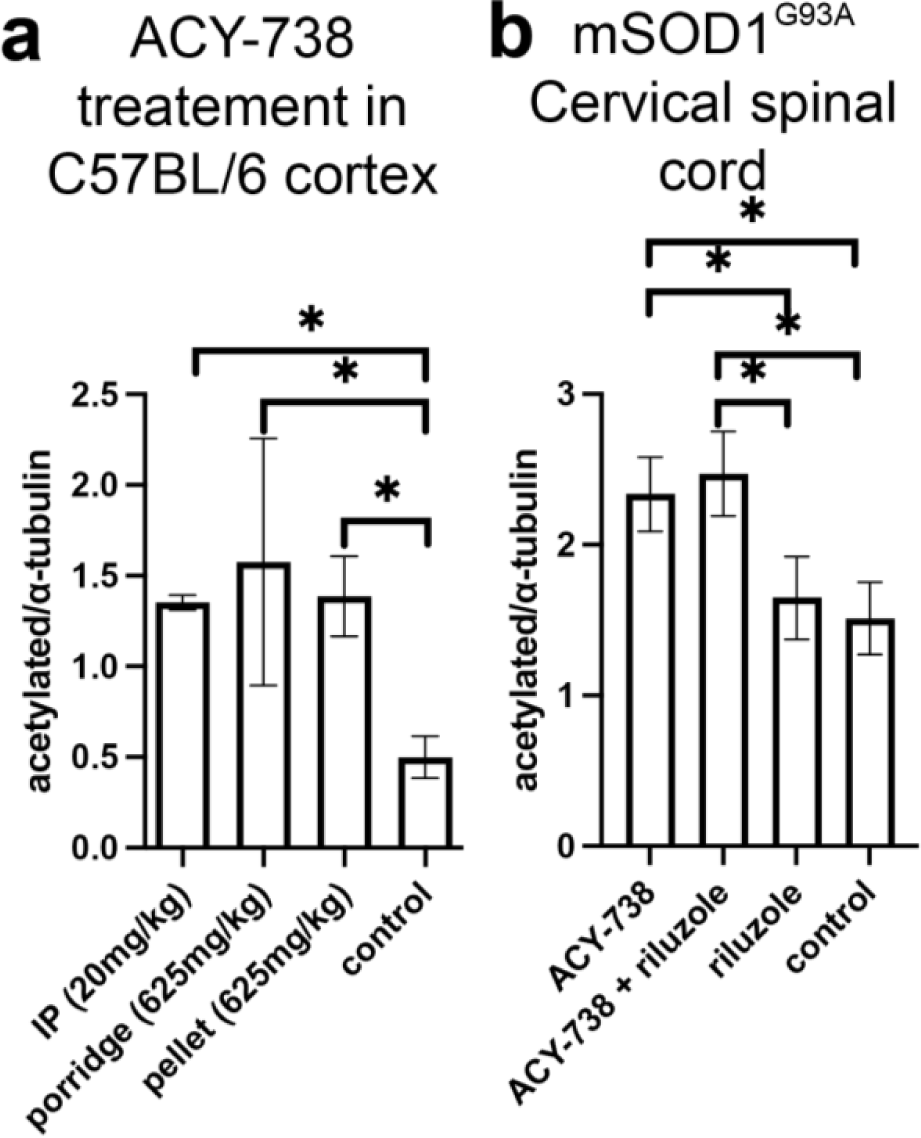
ACY-738 inhibits HDAC6 activity in the spinal cord of mSOD1^G93A^ mice. Indirect ELISA of the ratio of acetylated α-tubulin to α-tubulin show acute treatment of ACY-738 for 3 days in C57BL/6 mice led to increased cortical acetylated α-tubulin (a). Treatment administered by intraperitoneal (IP) injection (20mg/kg bodyweight), oral dosing via porridge (625mg/kg chow) or formulated into pellets (625mg/kg chow) did not significantly alter acetylated α-tubulin levels (a). Bar graph of ratios for relative indirect ELISA absorbance show treatment with ACY-738, and co-dosing ACY-738+riluzole increased acetylated α-tubulin in 20-week-old mSOD1^G93A^ mice compared to riluzole only, and untreated control groups (b). Data presented represent mean ± 95% confidence intervals. N=5-8 per treatment group.

**Figure 2:**
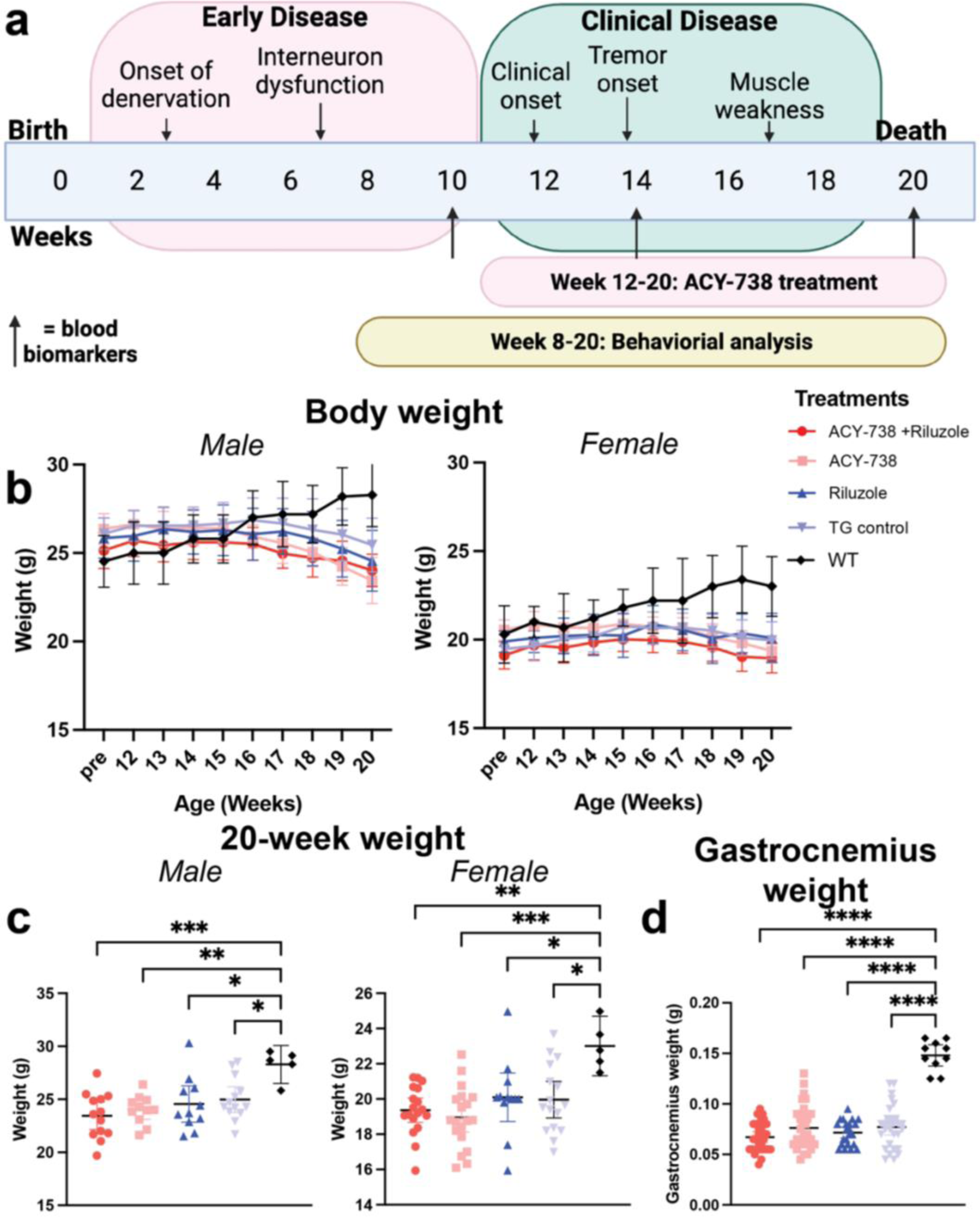
HDAC6i does not prevent weight loss and muscle deterioration in mSOD1^G93A^ mice. Schematic illustration of clinicopathological features of the mSOD1^G93A^ mouse model and experimental timeline for behavioural testing and endpoint (a). Body weight is unaltered by chronic treatment with ACY-738 (salmon), ACY-738 + riluzole (red), riluzole (blue) in mSOD1^G93A^ mice compared to untreated mSOD1^G93A^ (light blue) controls, however, are significantly lower than C57BL/6 controls (black) at endpoint (b). Treatment with ACY-738 did not rescue disease-associated weight loss in male (c - left), or female (c – right) mSOD1^G93A^ mice at 20 weeks of age. Gastrocnemius weight is significantly decreased in pooled male and female mSOD1^G93A^ mice compared to C57BL/6 control group, regardless of treatment with ACY-738 or riluzole (d). Data presented represent mean ± 95% CI. N=5 for C57BL/6 mice and n=11-19 per mSOD1^G93A^ treatment group.

Some of the earliest changes in ALS include the decline in motor function, which is recapitulated in the mSOD1^G93A^ mice (Gurney et al., 1994). To determine whether ACY-738 delayed the onset of motor symptoms in mSOD1^G93A^ mice, motor function tests were performed weekly until late-stage disease, including grip strength, wire-hang, and NeuroScore (Hatzipetros et al., 2015) tests. Gross motor strength was measured as all paw grip strength from 9-20 weeks of age. Initially, statistical analysis was performed comparing untreated mSOD1^G93A^ mice to untreated C57BL/6 mice. As expected, there was a significant genotype by age interaction, where untreated mSOD1^G93A^ mice exhibited significantly lower grip strength, that worsened over time compared to untreated C57BL/6 controls (F_1,1284_ =632.44, p <0.05, ANOVA type III Satterthwaite’s; Table 2; Figure 3 a-b). Comparisons of mSOD1^G93A^ mice treated with ACY-738, riluzole, ACY-738+riluzole and untreated mSOD1^G93A^ animals showed two significant 3-way interactions. There was a significant 3-way interaction between ACY-738, sex and age (F_1,1277_ =6.29, p<0.05, ANOVA type III Satterthwaite’s; Table 2; Figure 3 a-b) where ACY-738 treatment worsened grip strength in male mSOD1^G93A^ mice compared to untreated male mice. There was also a significant 3-way interaction between riluzole, sex, and age (F_1,1277_ =6.39, p<0.05, ANOVA type III Satterthwaite’s; Table 2; Figure 3 a-b) with riluzole treated mSOD1^G93A^ female mice losing grip strength quicker than untreated mSOD1^G93A^ controls (Figure 3 c). Together these data indicated that ACY-738 and riluzole both independently altered all-paw grip strength in mSOD1^G93A^ mice, with ACY-738 reducing grip-strength in male, but not female, mSOD1^G93A^ mice.

**Table 2:** All paw grip strength comparison between mSOD1^G93A^ and C57BL/6 control mice.

| *Predictors* | *Sex* | *N* | *mean* | *95%CI* |
| --- | --- | --- | --- | --- |
| C57BL/6 | F | 5 | 211 | 191 – 231 |
| C57BL/6 | M | 5 | 216 | 188 - 245 |
| mSOD1<sup>G93A</sup> vehicle | F | 16 | 118 | 101 - 136 |
| mSOD1<sup>G93A</sup> vehicle | M | 15 | 130 | 112 - 148 |
| mSOD1<sup>G93A</sup> riluzole | F | 12 | 129 | 109 - 149 |
| mSOD1<sup>G93A</sup> riluzole | M | 11 | 141 | 120 - 162 |
| mSOD1<sup>G93A</sup> ACY-738 | F | 18 | 117 | 101 - 134 |
| mSOD1<sup>G93A</sup> ACY-738 | M | 15 | 124 | 107 - 142 |
| mSOD1<sup>G93A</sup> ACY-738 + riluzole | F | 20 | 123 | 108 - 139 |
| mSOD1<sup>G93A</sup> ACY-738 + riluzole | M | 11 | 135 | 114 - 155 |

*Data presented as conditional means and 95% confidence intervals.*

The wire-hang task measures muscle performance, coordination, and endurance in rodent models (Rogers et al., 1997). mSOD1^G93A^ mice and C57BL/6 controls performed the hanging wire task at 12 weeks (treatment onset), 16 weeks, and 20 weeks (late-stage disease) of age. Initial analysis of untreated mSOD1^G93A^ mice and untreated C57BL/6 controls showed a significant main effect of genotype (F_1,37_ =47, p<0.05, ANOVA type III Satterthwaite’s; Table 3; Figure 3), and age (F_1,80_ =4.66, p = <0.05, ANOVA type III Satterthwaite’s; Table 3; Figure 3), but no significant effect of sex, where mSOD1^G93A^ mice fell sooner than C57BL/6 control controls, which occurred sooner, as disease progressed. There was a significant effect of age (F_1,249_ =48.83, p<0.05, ANOVA type III Satterthwaite’s; Table 3; Figure 3), and sex (F_1,123_ =14.60, p<0.05, ANOVA type III Satterthwaite’s; Table 3; Figure 3), on the latency to fall for treated mSOD1^G93A^ mice, whereby all male mSOD1^G93A^ mice fell quicker than female mice, regardless of treatment group, and latency to fall reduced with disease progression, regardless of mSOD1^G93A^ treatment group. However, there was no significant main effects or interactions with ACY-738 or riluzole on the wire-hang task for mSOD^G93A^ mice (Table 3). Together, these data indicated that ACY-738 and riluzole did not improve motor performance on wire-hang tasks in mSOD^G93A^ mice.

**Table 3:** Latency to fall from wire-hang task is not altered after treatment with ACY-738.

| *Predictors* | *Sex* | *N* | *Mean (s)* | *95%CI* |
| --- | --- | --- | --- | --- |
| WT | F | 5 | 136.7 | 103.1 – 170.3 |
| WT | M | 5 | 157.7 | 120.8 – 194.5 |
| mSOD1<sup>G93A</sup> vehicle | F | 16 | 3.67 | 3.32 - 4.01 |
| mSOD1<sup>G93A</sup> vehicle | M | 15 | 3.68 | 3.33 – 4.04 |
| mSOD1<sup>G93A</sup> riluzole | F | 14 | 3.93 | 3.49 - 4.37 |
| mSOD1<sup>G93A</sup> riluzole | M | 11 | 3.07 | 2.59 - 3.54 |
| mSOD1<sup>G93A</sup> ACY-738 | F | 18 | 3.81 | 3.44 – 4.18 |
| mSOD1<sup>G93A</sup> ACY-738 | M | 15 | 3.02 | 2.59 - 3.54 |
| mSOD1<sup>G93A</sup> ACY-738 + riluzole | F | 19 | 3.85 | 3.49 – 4.21 |
| mSOD1<sup>G93A</sup> ACY-738 + riluzole | M | 11 | 2.98 | 2.50 – 3.46 |

*Data presented as conditional means and 95% confidence intervals.*

As part of the NeuroScore paradigm, the development of muscle tremor, another measure of worsening motor system decline and muscle degeneration, was investigated in mSOD1^G93A^ mice (Gurney et al., 1994). While no tremor was observed for C57BL/6 mice at any age, mSOD1^G93A^ mice developed muscle tremors as the disease progressed, where most mSOD1^G93A^ mice had developed tremors by week 15. The week of tremor onset was measured, and Kaplan-Meier time-to-event analysis showed a significant effect of ACY-738 treatment on tremor onset in mSOD1^G93A^ mice, where ACY-738 treatment led to an earlier onset of tremor at 13 weeks of age compared to untreated mSOD1^G93A^ controls (p<0.05, Kaplan-Meier; Table 4; Figure 3). There was no significant effect of riluzole treatment on tremor onset in the mSOD1^G93A^ mice, with or without ACY738 administration (F_1,123_ = 9.42, p<0.05, Kaplan-Meier; Table 4; Figure 3). Together, these data indicated that ACY-738 treatment may detrimentally contribute to motor function.

**Table 4:** Neuromuscular tremor onset in mSOD1^G93A^ mice treated with ACY-738.

| *Predictors* | *Sex* | *N* | *Mean (week of onset)* | *95%CI* |
| --- | --- | --- | --- | --- |
| mSOD1<sup>G93A</sup> vehicle | F | 16 | 15.6 | 14.8 – 16.5 |
| mSOD1<sup>G93A</sup> vehicle | M | 14 | 15.0 | 14.1 – 16.0 |
| mSOD1<sup>G93A</sup> riluzole | F | 12 | 15.7 | 14.7 – 16.8 |
| mSOD1<sup>G93A</sup> riluzole | M | 11 | 15.1 | 14.1 – 16.2 |
| mSOD1<sup>G93A</sup> ACY-738 | F | 18 | 14.7 | 13.9 – 15.6 |
| mSOD1<sup>G93A</sup> ACY-738 | M | 15 | 13.9 | 13.0 – 14.9 |
| mSOD1<sup>G93A</sup> ACY-738 + riluzole | F | 20 | 14.9 | 14.1 – 15.7 |
| mSOD1<sup>G93A</sup> ACY-738 + riluzole | M | 11 | 14.1 | 13.1 – 15.1 |

*Data presented as conditional means and 95% confidence intervals.*

**Figure 3:**
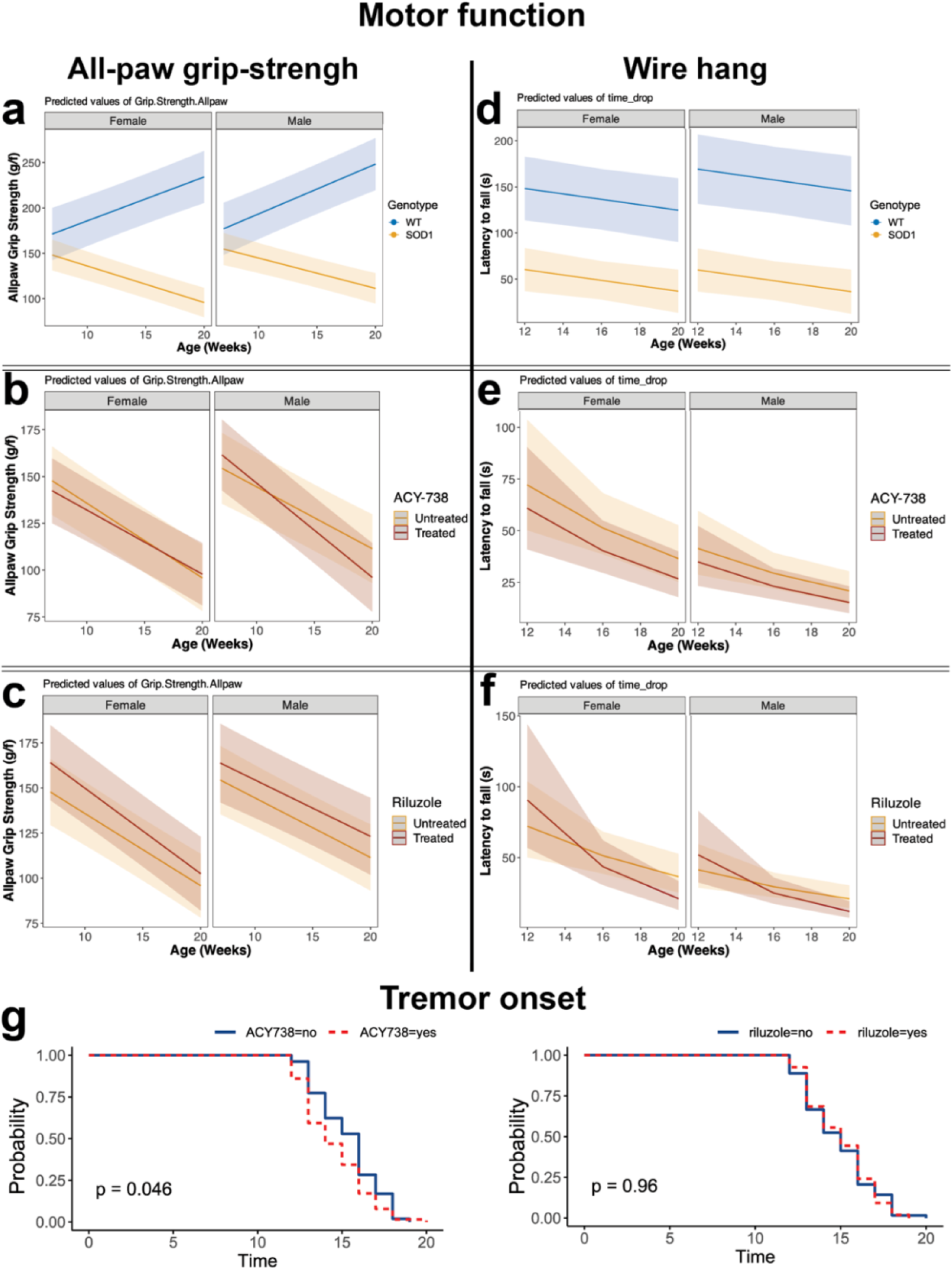
Motor function decline is unaltered by treatment with ACY-738 in mSOD1^G93A^ mice. Mixed effects models of all-paw grip-strength (a-c) for female (left) and male (right) mSOD1^G93A^ mice (yellow), compared to C57BL/6 (blue) control littermates, showed a significant reduction in all-paw grip-strength in mSOD1^G93A^ mice (a). Treatment with ACY-738 (b) or riluzole (c) (red) led to sex-specific alterations in all-paw grip strength in mSOD1^G93A^ animals. Wire-hang performance was significantly reduced in mSOD1^G93A^ (yellow) mice compared to C57BL/6 control littermates (blue) (d). Wire-hang was unaltered after treatment with ACY-738 (e), or riluzole (f) (red) in mSOD1^G93A^ mice. Kaplan-Meier time-to-event analysis demonstrates a significant effect of ACY-738 treatment on muscle tremor onset, compared to untreated mSOD1^G93A^ controls (g – left, ACY-738 = red), whereas riluzole treatment does not alter tremor onset in mSOD1^G93A^ mice (g – right, riluzole = red). Data presented represent conditional mean ± 95% CI for predicted values. N=11-19 per treatment group, and n=5 for C57BL/6 controls.

### Lower motor neurons in the lumbar spinal cord are rescued in female mSOD1 mice by ACY-738

Given that ACY-738 had mixed effects on motor performance in mSOD1^G93A^ mice, we examined whether ACY-738 rescued lumbar spinal cord motor neurons from degeneration (Gurney et al., 1994). Lower motor neurons from the lumbar spinal cord ventral horn were counted in cresyl violet stained sections taken at end point (20 weeks). Untreated mSOD1^G93A^ mice displayed reduced lower motor neuron numbers compared to C57BL/6 controls, indicating loss (F_1,456.71_ = 217, p <0.05, ANOVA type III Satterthwaite’s; Table 7; Figure 4b). When analysing mSOD1^G93A^ mice treated with ACY-738, riluzole, or co-treatment, there was a significant three-way interaction between ACY-738, riluzole and sex (p<0.05; gaussian mixed effects model; Type III Wald; Table 7; Figure 4). However, female mice treated with ACY-738 (p= 0.002) or riluzole (p= 0.013) alone had more lower motor neurons (figure 4c), indicating motor neurons were rescued by ACY-738 or riluzole treatment in these animals. Female mice co-treated with both drugs had motor neuron counts similar to baseline untreated mSOD1^G93A^ female mice (Figure 4). These data indicate that complex interactions may occur between ACY-738 treatment and riluzole, and disease onset and severity may need to be further considered for treatment paradigms.

### Axon puncta size is partially rescued in mice treated with ACY-738 and riluzole

Although ACY-738 treatment did not restore gross motor function, we sought to determine whether there were underlying changes in peripheral axons vulnerable to ALS pathology. Analysis of axonal dysfunction and degeneration in the sciatic nerve was measured with the pan-axonal marker against phosphorylated neurofilaments medium and heavy, SMI312 (Masliah et al., 1993). Measurements of SMI312 immunoreactivity within axons showed a significant reduction in the size of SMI312 positive axon puncta in mSOD1^G93A^ mice at 20 weeks of age, compared to C57BL/6 littermates (F_1,19.72_ = 13.88, p<0.05, ANOVA type III Satterthwaite’s; Table 5; Figure 4e). Male and female mSOD1^G93A^ mice had similarly sized SMI312 axonal puncta, indicating no significant difference between sexes.

There was no main effect of ACY-738 treatment or sex on SMI312 axon size in mSOD1^G93A^ treated mice, suggesting SMI312 immunoreactivity is consistent between male and female mSOD1^G93A^ mice. Furthermore, there were no interactions between ACY-738, riluzole treatment or sex. There was, however, a main effect of riluzole, with riluzole-treated mSOD1^G93A^ mice having significantly larger SMI312 positive axons than untreated mice (F_1,60.50_ = 4.02, p<0.05, ANOVA type III Satterthwaite’s; Table 5; Figure 4). There were no significant interactions or main effects for ACY-738, riluzole or sex on the number of SMI312 positive axons or the percentage area covered by SMI312 positive axons in mSOD1^G93A^ mice.

Choline Acetyltransferase, (ChAT) is a marker of motor neuron axons that are vulnerable to degeneration in ALS. ChAT labelling was utilised to determine if ACY-738 or riluzole treatment was altering motor axon puncta size specifically in peripheral nerves. Analysis of average axon size for ChAT-positive axons showed axon puncta size was significantly smaller in mSOD1^G93A^ mice compared to C57BL/6 controls (F_1,21.3_ = 8.43, p<0.05, ANOVA type III Satterthwaite’s; Table 5; Figure 4). Further analysis of mSOD1^G93A^ treated mice showed a significant 2-way interaction between ACY-738 and riluzole treatment (F_1,59.23_ = 4.19, p<0.05, ANOVA type III Satterthwaite’s; Table 6; Figure 4), where co-treatment yielded larger ChAT positive axons in male and female mice. However, post-hoc multiple comparisons between ACY and riluzole treated mSOD1^G93A^ mice were not significant; thus, the driving effect could not be determined. Together, these data demonstrated that ACY-738 and riluzole led to partial restoration of SMI-312 and ChAT positive axon size. It is important to note that the observed changes in axon size after treatment with ACY-738 or riluzole could reflect either change to the overall axon size, or underlying changes to ChAT and neurofilaments the axons.

**Figure 4:**
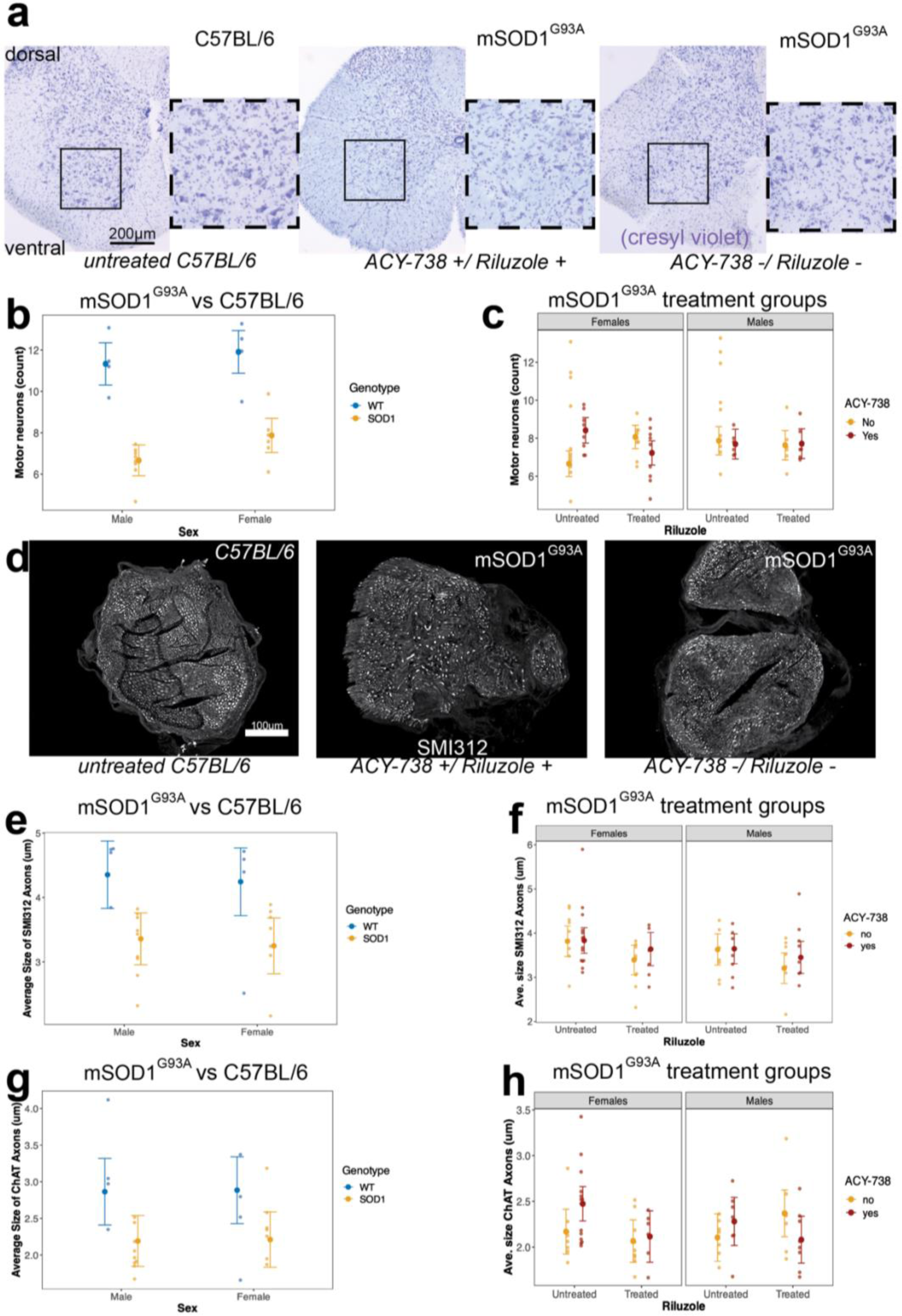
Motor neurons are rescued by ACY-738 treatment, and riluzole treatment improves axon fibres in mSOD1^G93A^ mice. Representative images of cresyl violet labelled lumbar spinal cord from ACY-738 and riluzole treated (left), vehicle treated mSOD1^G93A^ mice (middle), and C57BL/6 vehicle controls (right) (a). Motor neurons are significantly reduced in the lumbar spinal cord of mSOD1^G93A^ mice compared to C57BL/6 controls (b). Lower motor neurons are rescued by treatment with riluzole or ACY-738 individually, but not co-treatment in female mSOD1^G93A^ mice (c). Representative images of sciatic nerve from ACY-738 and riluzole treated (left), vehicle treated mSOD1^G93A^ mice (middle), and C57BL/6 vehicle controls (right) (d). The average size of SMI312 positive axons in the sciatic nerve are significantly reduced in mSOD1^G93A^ mice compared to C57BL/6 control littermates (e). Riluzole restores the average size of SMI312 positive axons in the sciatic nerve of mSOD1^G93A^ mice (f). The average size of ChAT positive axons in the sciatic nerve are significantly reduced in mSOD1^G93A^ mice compared to C57BL/6 control littermates (g). Combined treatment of riluzole and ACY-738 restores the average size of ChAT positive axons in the sciatic nerve of mSOD1^G93A^ mice (h). Data presented as conditional means and 95% confidence intervals. n=6-9 per treatment group, and n=4 for C57BL/6 controls.

**Table 5:** SMI312 positive axons are rescued by riluzole treatment in mSOD1^G93A^ mice.

| *Predictors* | *Sex* | *N* | *mean* | *95%CI* |
| --- | --- | --- | --- | --- |
| C57BL/6 WT | F | 4 | 4.35 | 3.83 – 4.88 |
| C57BL/6 WT | M | 4 | 4.24 | 3.72 – 4.77 |
| mSOD1<sup>G93A</sup> vehicle | F | 9 | 3.39 | 3.06 – 3.73 |
| mSOD1<sup>G93A</sup> vehicle | M | 7 | 3.21 | 2.86 – 3.55 |
| mSOD1<sup>G93A</sup> riluzole | F | 8 | 3.82 | 3.47 – 4.16 |
| mSOD1<sup>G93A</sup> riluzole | M | 7 | 3.63 | 3.28 – 3.98 |
| mSOD1<sup>G93A</sup> ACY-738 | F | 6 | 3.64 | 3.26 – 4.01 |
| mSOD1<sup>G93A</sup> ACY-738 | M | 8 | 3.45 | 3.09 – 3.82 |
| mSOD1<sup>G93A</sup> ACY-738 + riluzole | F | 14 | 3.83 | 3.54 – 4.12 |
| mSOD1<sup>G93A</sup> ACY-738 + riluzole | M | 7 | 3.65 | 3.31 – 3.98 |

*Data presented as conditional means and 95% confidence intervals.*

**Table 6:** ChAT positive axons are rescued by ACY-738 and riluzole treatment in female mSOD1^G93A^ mice.

| *Predictors* | *Sex* | *N* | *mean* | *95%CI* |
| --- | --- | --- | --- | --- |
| C57BL/6 WT | F | 4 | 2.86 | 2.41 – 3.32 |
| C57BL/6 WT | M | 4 | 2.88 | 2.43 – 3.34 |
| mSOD1<sup>G93A</sup> vehicle | F | 9 | 2.07 | 1.83 – 2.30 |
| mSOD1<sup>G93A</sup> vehicle | M | 7 | 2.37 | 2.11 – 2.62 |
| mSOD1<sup>G93A</sup> riluzole | F | 8 | 2.17 | 1.92 – 2.41 |
| mSOD1<sup>G93A</sup> riluzole | M | 7 | 2.10 | 1.84 – 2.36 |
| mSOD1<sup>G93A</sup> ACY-738 | F | 6 | 2.12 | 1.83 – 2.40 |
| mSOD1<sup>G93A</sup> ACY-738 | M | 8 | 2.08 | 1.82 – 2.34 |
| mSOD1<sup>G93A</sup> ACY-738 + riluzole | F | 14 | 2.47 | 2.29 – 2.66 |
| mSOD1<sup>G93A</sup> ACY-738 + riluzole | M | 7 | 2.28 | 2.02 – 2.54 |

*Data presented as conditional means and 95% confidence intervals.*

**Table 7:** Lumbar motor neuron counts in mSOD1^G93A^ mice treated with ACY-738.

| *Predictors* | *Sex* | *N* | *mean* | *95%CI* |
| --- | --- | --- | --- | --- |
| C57BL/6 WT | F | 4 | 11.33 | 10.31 – 12.34 |
| C57BL/6 WT | M | 4 | 11.90 | 10.88 – 12.93 |
| mSOD1<sup>G93A</sup> vehicle | F | 8 | 6.66 | 5.98 – 7.33 |
| mSOD1<sup>G93A</sup> vehicle | M | 6 | 7.86 | 7.12 – 8.61 |
| mSOD1<sup>G93A</sup> riluzole | F | 9 | 8.07 | 7.45 – 8.68 |
| mSOD1<sup>G93A</sup> riluzole | M | 6 | 7.63 | 6.86 – 8.40 |
| mSOD1<sup>G93A</sup> ACY-738 | F | 8 | 8.41 | 7.74 – 9.08 |
| mSOD1<sup>G93A</sup> ACY-738 | M | 6 | 7.69 | 6.90 – 8.47 |
| mSOD1<sup>G93A</sup> ACY-738 + riluzole | F | 9 | 7.23 | 6.59 – 7.86 |
| mSOD1<sup>G93A</sup> ACY-738 + riluzole | M | 6 | 7.71 | 6.93 – 8.49 |

*Data presented as conditional means and 95% confidence intervals.*

### Iba1 positive microglia are unaltered after treatment with ACY-738 in mSOD1^G93A^ mice

Glial reactivity in the spinal cord is a common hallmark of ALS. To determine if ACY-738 treatment lowered glial reactivity in the lumbar spinal cord, Iba1 positive microglia were immunolabelled, imaged, and quantified as a percentage area of the ventral horns. Initially, when comparing untreated mSOD1^G93A^ and C57BL/6 mice, there was a significant main effect of genotype (F_1,25.94_ = 19.40, p<0.05, ANOVA type III Satterthwaite’s; Table 8; Figure 5), whereby mSOD1^G93A^ mice had a higher percentage area of Iba1 positive staining compared to C57B/L6 mice. However, there were no significant interactions between genotype or sex, and no main effect of sex on the percentage area of Iba1 positive microglia in mSOD1^G93A^ mice. A comparison of mSOD1^G93A^ mice treated with ACY-738, riluzole, or co-treatment with ACY-738 and riluzole showed no significant main effects of ACY-738 or riluzole treatment on the percentage area covered by Iba1 immunoreactivity (Table 8). These data suggest that ACY-738 does not alter reactive gliosis in the spinal cord of mSOD1^G93A^ mice.

**Table 8:** Iba1 positive microglia are unaltered after treatment with ACY-738.

| *Predictors* | *Sex* | *N* | *mean* | *95%CI* |
| --- | --- | --- | --- | --- |
| C57BL/6 WT | F | 4 | 3.75 | -0.07 – 7.58 |
| C57BL/6 WT | M | 4 | 2.26 | -1.57 – 6.09 |
| mSOD1<sup>G93A</sup> vehicle | F | 11 | 12.18 | 9.60 – 15.0 |
| mSOD1<sup>G93A</sup> vehicle | M | 10 | 11.75 | 7.99 – 13.6 |
| mSOD1<sup>G93A</sup> riluzole | F | 8 | 13.45 | 8.53 – 18.4 |
| mSOD1<sup>G93A</sup> riluzole | M | 7 | 13.23 | 7.98 – 18.5 |
| mSOD1<sup>G93A</sup> ACY-738 | F | 11 | 12.10 | 7.90 – 16.3 |
| mSOD1<sup>G93A</sup> ACY-738 | M | 10 | 10.26 | 5.87 – 14.7 |
| mSOD1<sup>G93A</sup> ACY-738 + riluzole | F | 16 | 9.04 | 5.57 – 12.5 |
| mSOD1<sup>G93A</sup> ACY-738 + riluzole | M | 8 | 9.62 | 4.69 – 14.6 |

*Data presented as conditional means and 95% confidence intervals.*

### GFAP labelled astrocytes are reduced in the lumbar spinal cord after treatment with riluzole, but not ACY-738 in mSOD1^G93A^ mice

Astrocyte reactivity is greatly increased in mSOD1^G93A^ mice, coinciding with pathology accumulation and lower motor neuron loss (Yamanaka & Komine, 2018). GFAP immunoreactivity increased in the ventral horn of the lumber spinal cord in mSOD1^G93A^ mice at end point compared to C57BL/6 WT controls, demonstrating a significant main effect of genotype in the percentage area of GFAP positive astrocytes (F_1,24.022_ = 35.04, p<0.05, ANOVA type III Satterthwaite’s; Table 9; Figure 5). Untreated male and female mSOD1^G93A^ and C57BL/6 mice, had a similar percentage area of GFAP positive ventral horn astrocytes, highlighting no significant interactions between genotype or sex, and no main effect of sex in these animals. Treatment of ACY-738 with or without riluzole, did not reduce the percentage area of GFAP positive astrocytes, whilst riluzole alone reduced the percentage area of GFAP positive labelling in mSOD1^G93A^ mice, compared to untreated mSOD1^G93A^ mice (Figure 5). These results showed a significant main effect of riluzole treatment on GFAP labelled astrocytes (F_1,59.80_ = 4.38, p<0.05, ANOVA type III Satterthwaite’s; Table 9; Figure 5), but no main effect of ACY-738 treatment or interactions between ACY-738 and riluzole in the spinal cord ventral horn of mSOD1^G93A^ mice. There was also no main effect of sex in the ventral horn of mSOD1^G93A^ mice treated with ACY-738, riluzole, or co-treatment. These data suggest that riluzole, but not ACY-738 treatment, reduce reactive astrogliosis in the ventral horn of the lumbar spinal cord in mSOD1^G93A^ mice.

**Table 9:** Astrocyte reactivity is unaltered in mSOD1^G93A^ mice treated with ACY-738.

| *Predictors* | *Sex* | *N* | *mean* | *95%CI* |
| --- | --- | --- | --- | --- |
| C57BL/6 WT | F | 4 | 1.92 | -1.55 – 5.38 |
| C57BL/6 WT | M | 4 | 0.44 | -3.04 – 3.92 |
| mSOD1<sup>G93A</sup> vehicle | F | 11 | 12.72 | 9.70 – 15.7 |
| mSOD1<sup>G93A</sup> vehicle | M | 7 | 10.30 | 6.56 – 14.0 |
| mSOD1<sup>G93A</sup> riluzole | F | 8 | 9.07 | 5.55 – 12.6 |
| mSOD1<sup>G93A</sup> riluzole | M | 7 | 11.74 | 8.01 – 15.5 |
| mSOD1<sup>G93A</sup> ACY-738 | F | 9 | 16.44 | 13.12 – 19.8 |
| mSOD1<sup>G93A</sup> ACY-738 | M | 10 | 12.84 | 9.71 – 16.0 |
| mSOD1<sup>G93A</sup> ACY-738 + riluzole | F | 14 | 12.08 | 9.45 – 14.7 |
| mSOD1<sup>G93A</sup> ACY-738 + riluzole | M | 8 | 9.07 | 6.01 – 13.0 |

**Figure 5:**
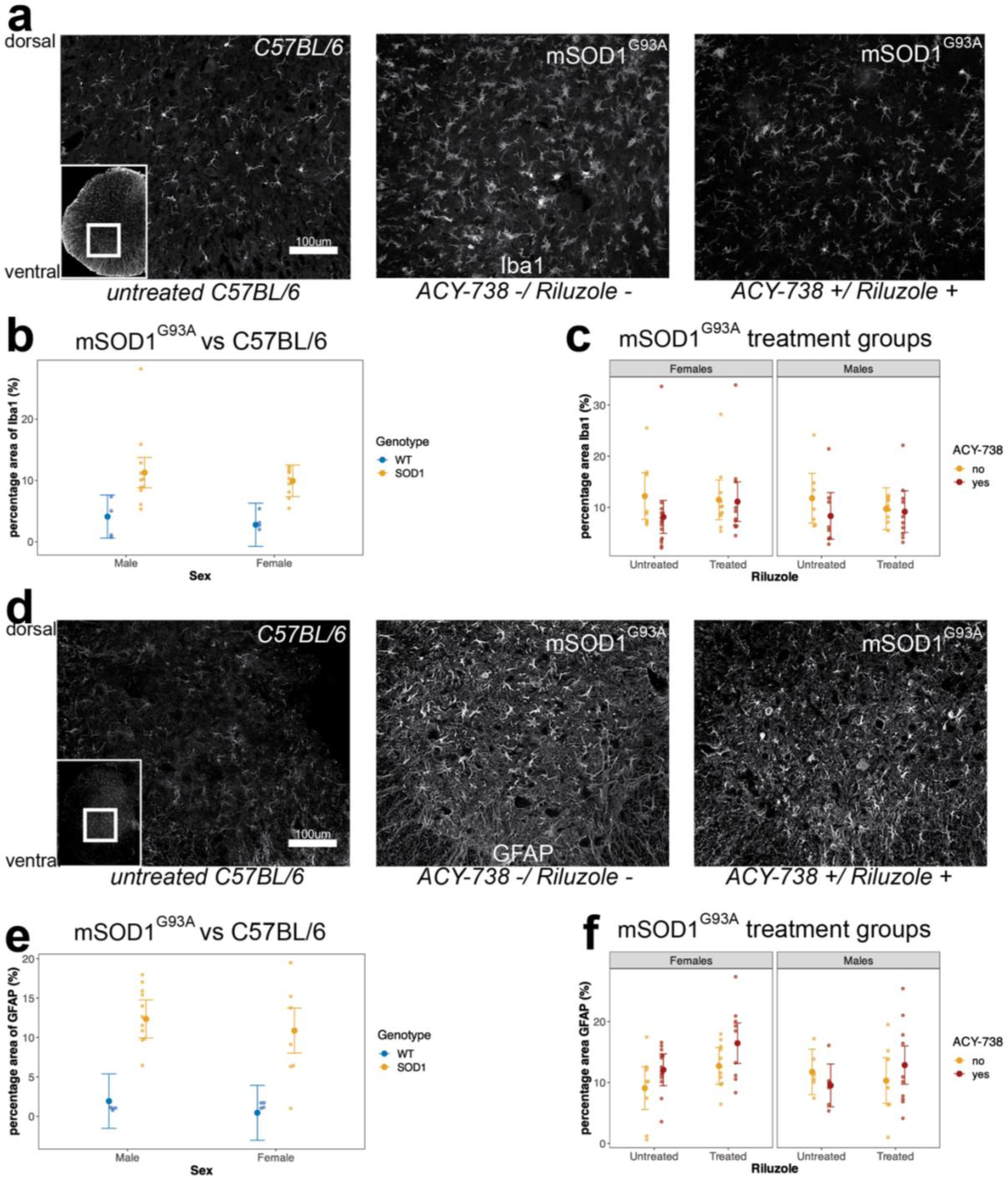
Astrocytes are altered after treatment with riluzole and ACY-738 in mSOD1^G93A^ mice. Representative images of Iba1+ve microglia in the lumbar spinal cord of mSOD1^G93A^ mice treated with ACY-738 (left), untreated (middle), and C57BL/6 wildtype (right) mice (a). Scatter plot showing mSOD1^G93A^ mice (right) have significantly increased microglia in the lumbar spinal cord compared to C57BL/6 WT (left) control littermates (b). Scatter plot showing mSOD1^G93A^ microglia in the lumbar spinal cord are unaltered after treatment with ACY-738, riluzole, or both (c). Representative images of GFAP positive astrocytes in the lumbar spinal cord of mSOD1^G93A^ mice treated with ACY-738 (left), untreated (middle), and C57BL/6 wildtype (right) mice (d). mSOD1^G93A^ mice have significantly increased GFAP astrocyte reactivity in the lumbar spinal cord compared to C57BL/6 WT controls (e). Female mSOD1^G93A^ mice treated with ACY-738 alone have significantly enriched GFAP reactivity compared to untreated littermates (f). Data presented as conditional means and 95% confidence intervals. N=5-9 per treatment group, and n=5 for C57BL/6 controls.

### Neurofilament light increases with disease progression but is not rescued by ACY-738 treatment in mSOD1^G93A^ mice

Alongside behavioural analyses, serum was collected at 14 and 20 weeks of age in mSOD1^G93A^ mice and WT controls for blood biomarker analysis. Analysis of serum was performed utilising Single Molecule Array (SIMOA) to measure neurofilament light (NFL), a biomarker of neuroaxonal degeneration. At 14 weeks, results demonstrated a significant effect of age (estimate = 1.43, CI=1.11-1.72, p<0.05; linear mixed effects model, Satterthwaite’s method; Supplementary Table 1), where there was an increase in NFL levels at 20 weeks compared to 14 weeks in mSOD1^G93A^ mice. There was also a significant effect of sex (estimate = 0.15, CI=-0.28-0.58, p<0.05; linear mixed effects model, Satterthwaite’s method; Supp. Table 1) on serum neurofilament levels in mSOD1^G93A^ mice, where males and females had similar levels of NFL at 14 weeks, but males had significantly higher NFL than females at 20 weeks. However, there was no observed effect of treatment with ACY-738 or riluzole on serum NFL levels (Figure 6). These results demonstrate that ACY-738 does not lead to detectible changes in neurofilament blood biomarkers in mSOD1^G93A^ mice.

**Figure 6:**
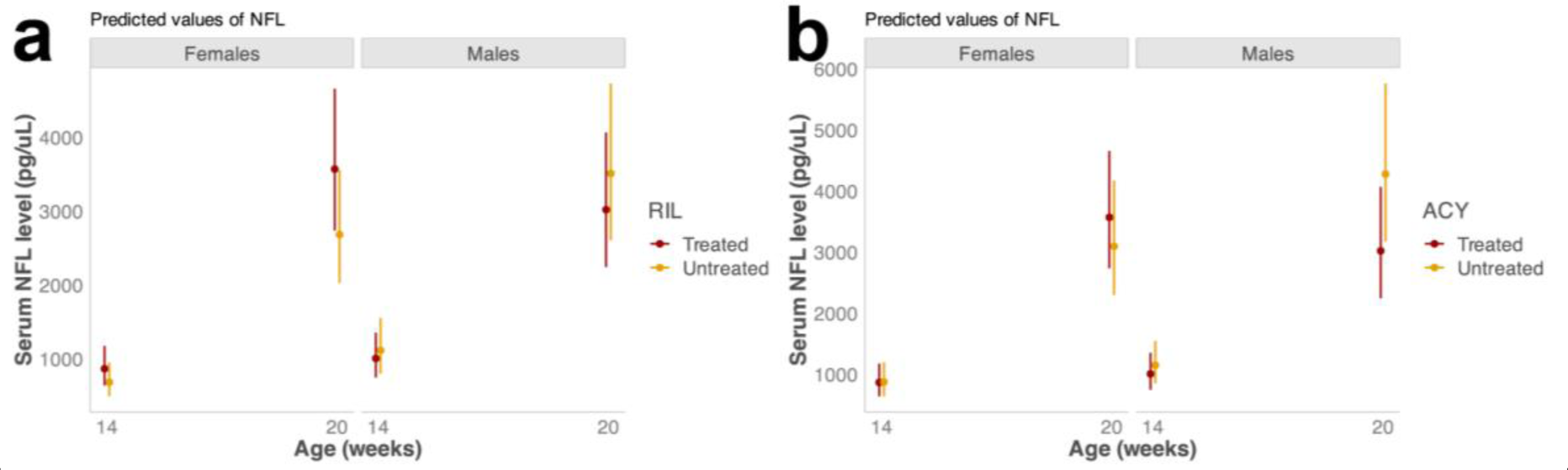
Serum NFL is significantly increased with disease progression but is unaltered by treatment with ACY-738 or riluzole. Plots showing predicted values of NFL in mSOD1^G93A^ mice treated with riluzole (a), or ACY-738 (b), at 14, and 20 weeks of age respectively. There was a significant increase in the level of serum NFL (log pg/uL) between 14 and 20 weeks of age. Data presented as conditional means and 95% confidence intervals.

## Discussion

In ALS, motor neuron axons are selectively vulnerable to degeneration, potentially through mechanisms independent of the cell soma (Gould et al., 2006), thus therapeutics targeting axon protection are emerging as novel treatments. The current study administered an HDAC6 inhibitor, ACY-738, to prevent the deacetylation of microtubules in the mSOD1^G93A^ mouse model of ALS. Chronic ACY-738 treatment had complex effects on motor function and ALS linked pathology in mSOD1 mice, worsening motor outcomes in male, but not female, mSOD1 mice; providing modest protection of ventral horn motor neurons, and in isolation, did not protect against a reduction in axon puncta size or lessen glial immunoreactivity in mSOD1 animals. Co-treatment of ACY-738 with riluzole, maintained axonal puncta size in mSOD1 mice, but offered no further protection of ALS-linked pathology or improved motor function.

Novel approaches using HDACi to stabilize axons in models of neurodegeneration, show promising results *in vitro* and *in vivo* (Chuang et al., 2009; Hanson et al., 2018; Sugai et al., 2004). The specific inhibition of HDAC6 by novel HDAC6i drugs including ACY-738, have provided degrees of neuroprotective effects in models of MS, CMT and FUS-ALS (Benoy et al., 2017; LoPresti, 2019; Majid et al.; Rossaert et al., 2019). We affirmed that ACY-738 passes the BBB and increases α-tubulin acetylation in the spinal cord of mSOD1^G93A^ and C57BL/6 mice (Benoy et al., 2017; Rossaert et al., 2019), however, chronic ACY-738 administration offered only modest, selective improvements in ALS-linked pathology and symptoms in mSOD1^G93A^ animals. Pan-HDACi administered in mSOD1^G93A^ animals improved survival, gait and motor function (Ryu et al., 2005), or delayed motor and muscle function deficits, but not survival (C. Rouaux et al., 2007), highlighting the complex relationship between HDACi and ALS pathology. However, the selective mechanism of action of HDACi and ACY-738 in particular, could be due to the ALS model system, as ACY-738 (100mg/kg) treatment prevented weight loss, improved grip strength, and prolonged survival in FUS ALS mice (Rossaert et al., 2019). Given ACY-738 administration failed to offer any real functional or pathological improvements in mSOD1 mice, our conflicting results may be reflective of inherent genetic and pathological differences between FUS and mSOD1 mouse models, such as different pathological accumulations, or disease durations. Thus, future experiments into the effectiveness of HDACi, particularly ACY-738, should take into consideration the ALS model system.

Motor deficits, a hallmark symptom of clinical and animal models of ALS, arise due to the degeneration and death of motor neurons and their surrounding circuits. Thus, preserving neurons within motor circuits, and maintaining axonal size are essential to synaptic activity, molecular homeostasis, and normal function of the nervous system (Perrot et al., 2007; Sainio et al., 2021). Large axons are vulnerable to degeneration in ALS (Lefebvre-Omar et al., 2023; Nguyen, Larivière, & Julien, 2000), thus, preserving axon size and caliber to maintain axonal function is of importance. Evidence in ALS models supporting the protection of motor neuron axons and soma after ACY-738 treatment is inconclusive. In FUS mice, treatment with 100mg/kg ACY-738 protected neuromuscular junctions, but did not rescue lower motor neurons (Rossaert et al., 2019), whilst 3mg/kg ACY-738 treatment in models of CMT improved motor nerve conductivity *in vitro* and neuromuscular junction innervation *in vivo* (Benoy et al., 2017). Furthermore, in patient-derived iPSC motor neurons containing FUS mutations, 1µM ACY-738 increased α-tubulin acetylation and restored axonal transport deficits for endoplasmic reticulum vesicles and mitochondria (Guo et al., 2017). In the current study, treatment with ACY-738 provided modest protection to lower motor neurons in mSOD1^G93A^ mice, and alone was unable to rescue/preserve peripheral axon degeneration in sciatic nerves. The current study also showed serum NFL, a blood biomarker of neurodegeneration, to be unaltered after treatment with ACY-738 or riluzole at 14 and 20 weeks of age, which may indicate that although there were subtle improvements in lower motor neurons and their axons after treatment with ACY-738, it may not be enough to overcome the severity of neurodegeneration seen in the mSOD1^G93A^ model. The contradictory evidence for a role of ACY-738 to protect motor neurons between FUS and mSOD1 models highlights the need for further research into ACY-738 and HDAC6i in a wider range of ALS models to determine its therapeutic potential.

In any therapeutic intervention, the administered drug dosage is an important consideration to limit off-target effects. In the current study, animals were dosed at 625mg/kg in chow, a dosage that was calculated from ACY-738 chow administration in APP/PS1 mice modelling Alzheimer’s disease, which led to ∼100mg/kg ACY-738 ingested (Majid et al., 2015). In these mice, ACY-738 at 625mg/kg showed improvements in axonal transport, and recovery of short-term memory and learning, demonstrating the potential for ACY-738 to be used in ALS-linked degenerative models. In a more recent study, FUS animals dosed with 100mg/kg ACY-738 in chow showed global HDACi, where ACY-738 exerted non-HDAC6 specific inhibition, which rescued motor deficits, improved survival and rescued axon-trafficking in FUS ALS mice. Rossaert and colleagues (2019) also demonstrated that at 100mg/kg, ACY-738 altered class 1 HDAC activity, and improved survival in HDAC6 knock out mice, implicating global changes to histone modifications to rescue ALS (Rossaert et al., 2019). Indeed, global HDACi may have more beneficial effects in ALS. Recently, combined treatment with global HDACi, sodium phenylbutyrate and taurursodial slowed functional decline in phase II clinical trials for ALS, and were subsequently given FDA approval to treat ALS (Paganoni et al., 2020; Sun, Li, & Bedlack, 2023). It is possible, that in the current study, a dosage of 625mg/kg may be ‘overstabilising’ axons, reducing their inherent flexibility and ability to adapt to different cues, and thus emerges as an inappropriate dose for selective peripheral axon protection in ALS models. Recent literature has implicated that the destabilisation of microtubules may be neuroprotective in TDP-43 models of ALS (Baughn et al., 2023), which highlights the need for more finely tuned therapeutics and the complex role of microtubules in axon transport and degeneration. Additionally, HDAC6 inhibition may have a range of different off-target effects in the cytoplasm of cells that are being altered due to ACY-738 treatment. HDAC6 is thought to interact with polyubiquitin and have important roles in the aggresome-autophagy pathways, and proteosome formation (Yan, 2014), all of which are implicated in ALS disease pathogenesis, and therefore may be altered by ACY-738 administration (Yan, 2014). HDAC6 also has domains to anchor proteins to the cytoplasm, which may lead to altered protein homeostasis, when HDAC6 is inhibited (Yan, 2014). These mechanisms may be indicative of the alteration of more complex biochemical pathways due to ACY-738 treatment and warrant further investigation. Understanding the off-target effects of ACY-738 at 625mg/kg is an important avenue of future research to selectively identify ACY-738 as an appropriate novel target for ALS-linked degeneration.

Unlike many preclinical drug trials in ALS, the current study incorporated riluzole co-treatment with ACY-738, to determine ACY-738 - riluzole drug interactions, and to establish if ACY-738 treatment alone would be more neuroprotective than current FDA approved treatments. Riluzole’s mechanism of action is thought to involve changes in glutamatergic neurotransmission and reducing the effects of excitotoxicity, a common pathological mechanism underlying ALS (Bellingham, 2011; Doble, 1996b; Lazarevic et al., 2018). While much of the literature surrounding riluzole in ALS has suggested a modest, but significant improvement in animal model and patient outcomes, the extent to which these changes alter axon degeneration in particular, is less studied (Bellingham, 2011). Riluzole (22mg/kg) treatment alone was ineffective in improving motor function or lifespan in mSOD1^G93A^, FUS and TDP-43 mouse models (Hogg et al., 2018). In contrast, in our mSOD1 animals, riluzole alone improved motor outcomes during early-disease, improved motor neuron survival, reduced spinal cord astrocyte reactivity and restored SMI312-labelled axon size in the sciatic nerve. Co-treatment of riluzole alongside multiple novel compounds, including a cyclohexane analogue that reduced SOD1 aggregation; NU-9, or elacridar; a P-glycoprotein inhibitor, rather than riluzole alone, increased upper motor neurite outgrowth and branching *in vitro* and significantly improved survival in mSOD1^G93A^ mice, respectively (Genç et al., 2022; Jablonski et al., 2014). In the current study, co-treatment of ACY-738 and riluzole in mSOD1^G93A^ mice led to multifaceted findings; co-treatment maintained ChAT positive axon size in the sciatic nerve, yet led to increased lower motor neuron degeneration in the lumbar spinal cord. Co-treatment with ACY-738 and riluzole suggests that widespread axon alterations are occurring during ALS-linked degeneration and that ACY-738 and riluzole combination therapy may offer selective protection of these axons. These data suggest riluzole treatment may have more varied outcomes in the treatment of ALS, and also highlight the importance of preclinical testing in conjunction with currently available treatment paradigms.

This study also sought to determine if sex-specific differences were present after treatment with ACY-738 or riluzole. Studies of the mSOD1^G93A^ mouse model of ALS have reported inherent sex differences, where males have differential disease phenotype or time-course (Cacabelos et al., 2016; Hatzipetros et al., 2015). In line with this literature, this study demonstrated sex differences in tremor onset, motor function, grip-strength, NFL blood biomarker levels, alongside interactions between sex and ACY-738 or riluzole treatment in lumbar motor neuron counts for mSOD1^G93A^ mice. These reported differences may be due to differing disease progression rates between males and females in mSOD1^G93A^ mice. Sex differences have been widely reported in ALS literature, both in preclinical and clinical settings. Cacabelos et al., (2016) showed early sex differences in mitochondrial dysfunction and oxidative stress in the spinal cord of mSOD1^G93A^ mice (Cacabelos et al., 2016), and similarly, studies have reported differential response to preclinical drug treatments in these mice (Kaneb et al., 2011). Multiple studies have implicated sex differences in TDP-43 and C9orf72 animal models, and patients with C9orf72-linked ALS (Trojsi et al., 2019; Williams et al., 2013). Whilst sex differences have been widely reported in ALS, there have been few studies of blood biomarkers in ALS, and none to date have reported sex differences in ALS cases (Gaiottino et al., 2013; Loeffler et al., 2020; Verde et al., 2019) However, the sex differences in disease phenotype and severity, alongside improved sensitivity of the assay platform used herein may account for the sex-specific observations in the current study. Together, these findings demonstrate the importance of investigating sex specific differences in disease progression and response to treatment.

## Conclusion

In conclusion, the data presented in this study demonstrates a varied role of ACY-738 as a novel drug candidate for ALS. The compound prevented some lower motor neuron and peripheral axon degeneration, but suggests detrimental effects in combination with riluzole. The current study highlighted the complex interactions between disease severity in animals co-treated with ACY-738 and riluzole and demonstrates the importance of such study designs for the translatability of preclinical research into the clinic, the importance of interaction mechanisms, and need for further investigations into the role of microtubule acetylation in ALS. Overall, there were limited protective effects of HDAC6i with ACY-738 on neurodegeneration in mSOD1^G93A^ mice.

## Materials and Methods

### Animal housing, husbandry, and drug treatment

The study was approved by the Animal Ethics Committee of the University of Tasmania (Ethics approval number A0017895) and designed in accordance with the Australian Code of Practice for the Care and Use of Animals for Scientific Purposes (NHMRC, 2004).

mSOD1^G93A^ (B6.Cg-Tg(*SOD1*^G93A^)1Gur/J; Jax no. 004435; *m*SOD1^G93A^) mice maintained on a C57BL/6/J background were utilized for this study. Mice were housed in individually ventilated cages at room temperature on a 12-h light-dark cycle with *ad libitum* access to food and water. Genotypes were confirmed by ear-clip PCR at 3 weeks of age (Transnetyx, USA). Animals were split into 5 experimental groups: mSOD1^G93A^ ACY-738 treated; mSOD1^G93A^ treated with ACY-738 and riluzole, mSOD1^G93A^ riluzole treated, mSOD1^G93A^ untreated controls, and C57BL/6 wildtype controls. ACY-738 (University of Tasmania, Australia) was incorporated into chow at 625mg/kg, while control chow was untreated (Specialty Feeds, Australia), and riluzole hydrochloride was given at 188mg/L dissolved into drinking water. All mice were weighed and monitored weekly for overall health, and additionally assessed for motor function. All mice began motor function testing from 8 weeks of age, started on 625mg/kg ACY chow from 12 weeks of age and were 20 weeks of age at ethical endpoint for tissue collection.

### Motor function testing

Phenotypic neurological scoring was performed to evaluate disease progression in SOD1^G93A^ mice as previously described (Hatzipetros et al., 2015). Motor function was additionally investigated by the hanging wire test, as previously described (Rogers et al., 1997). Briefly, mice are placed in the centre of a wire, suspended between two columns, approximately 30cm high, for a total of 3 minutes. Mice are scored for number of falls, number of times a mouse reaches a column and time until first reach/drop. The hanging wire test was performed weekly until 20 weeks of age end-point. Grip strength was also investigated to evaluate disease progression, as previously described (Takeshita et al., 2017). Briefly, using a Chatillon® DFE II (Ametek) force measurement gauge with a 45° declined wire grid, mice were tested for fore paw and all paw grip strength, measured as peak tension (gram force) in triplicate, then averaged across trials (Takeshita et al., 2017). Motor function tests were compared between genotypes, treatment groups and sexes.

### ACY-738 bioavailability

To determine the bioavailability of ACY-738, mice were treated with intraperitoneal injection of ACY-738 (20mg/kg), or ACY-738 was mixed into a porridge of animal chow, or formulated into pellets (Specialty Feeds, WA, Australia) at 625mg/kg. C57BL/6 mice were injected daily, or given *ad libitum* access to chow, from 12 weeks of age and sacrificed 3 days after treatment for biochemical analysis with indirect ELISA.

### Blood collection and biomarker analysis

During motor function testing, blood samples were collected via peri-mandibular bleeds at week 14, and blood collected transcardially at time of perfusion (20 weeks of age). Blood was left at room temperature to clot for at least 30 minutes and then centrifuged at 3000 × g for 15 minutes at 4°C, with resulting serum moved to a fresh tube and the process repeated. After the second centrifugation the resulting serum was moved to a fresh tube and stored at -80°C until biomarker analysis. For biomarker analysis, serum samples were thawed at room temperature for 1 hour, vortexed for 10 seconds and then centrifuged at 10 000 × g for 5 minutes to remove particulate matter. Serum NFL analysis was performed using the Quanterix™ NF-Light™ Advantage SR-X Kit (Cat no. 103400) on the SR-X platform as per the manufacturer’s instructions. Calibrators and high and low control samples provided in the kit were run as per the manufacturer’s instructions in duplicate and samples were run in single and diluted 1/8 by adding 12.5µl to 87.5µl of sample diluent.

### Tissue collection and processing

Mice were anaesthetized with isoflurane (5% isoflurane and oxygen) to prevent discomfort, before being terminally anaesthetized with an overdose of sodium pentobarbitone (300mg/kg body weight, intraperitoneal) transcardially, and perfused with 4% (w/v) paraformaldehyde (PFA, Sigma Aldrich, St. Louis, MO, USA) in 0.01M phosphate-buffered saline (PBS, Sigma Aldrich, St. Louis, MO, USA). Brain, spinal cord and sciatic nerves which were dissected for later immunohistochemistry were post-fixed in 4% PFA overnight at 4°C and stored in PBS containing 0.02% sodium azide.

### Biochemical Analysis

Cervical spinal cord and cortex protein were extracted using a lysis buffer of RIPA buffer (Sigma; R0278), protease inhibitor (Roche; 11836153001) and phosphatase inhibitor cocktail (Roche; 4906837001) and quantified using a Pierce™ BCA protein assay (Thermo Scientific; 23225). Protein samples were analysed by indirect ELISA for acetylated α-tubulin and total α-tubulin. Briefly, 31.25ng of cervical spinal cord protein was added to 100µL bicarbonate buffer (8.4g/L NaHCO_3_, 36g/L Na_2_CO_3_, pH 9.5), and incubated on ELISA Nunc plates (Thermo Scientific; NUN442404) overnight at 4°C on an orbital shaker. Samples were then washed in ELISA wash buffer (0.01M PBS with 0.05% Tween 20; Sigma; P7949) for 4x 2 minutes on an orbital shaker. Samples were then blocked in 3% BSA (Thermo Scientific; A9418) for one hour at room temperature on an orbital shaker, before being incubated with acetylated tubulin (Sigma; T6793-.2ml), or alpha tubulin (Abcam; AB52866) at 1:500 for 90 minutes at room temperature. Samples were then washed 4x 2 minutes in ELISA wash buffer, then incubated with polyclonal mouse (DAKO; P0447), or rabbit (DAKO; P0448) HRP at 1:1000, in ELISA blocking solution, and incubated for 45minutes. Samples were washed 4x 2 minutes, then exposed with 100µL 1-step ultra TMB-ELISA substrate solution (Thermo Scientific; 34028) for 5 minutes, and the reaction stopped with 1M HCl. Plates were read on a TECAN Spark plate reader at 450nm.

### Immunohistochemistry

Spinal cord and sciatic nerve tissue processed for immunohistochemistry was mounted in OCT (Tissuetek Sakura; IA018) and cryosectioned (Leica CM1850UV) at 40µm. Spinal cord and sciatic nerve sections were stored in 0.01M PBS containing 0.02% sodium azide (Sigma; S2002) until processed for immunohistochemistry as previously described (Collins et al., 2022). Tissue was blocked using Dako serum free protein block (Dako; X090930-2) for 30 minutes before primary antibody incubation. Primary antibodies (detailed in Table 11) were incubated with free floating tissue in PBS-X (0.01M PBS and 0.5% Triton-X) overnight at 4°C. Sections were washed three times with 0.01M PBS and incubated with Alexa fluor secondary antibodies diluted in PBS-X. Samples were then washed 3x 10 minutes and then incubated with DAPI in 0.01M PBS, then mounted on slides with DAKO fluroescent mounting media (Agilent DAKO; S302380-2).

**Table 11:**
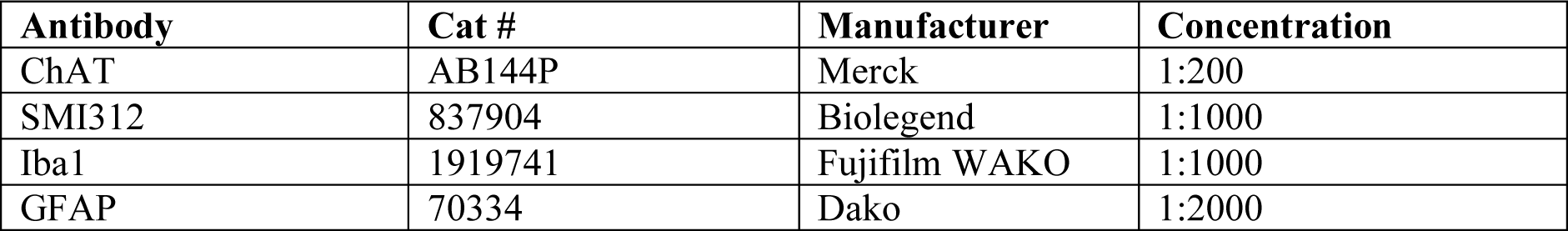
Primary antibodies.

### Image acquisition, quantitation, and analysis

Immunohistochemical images of spinal cord and sciatic nerve tissue were captured using an Olympus FV3000 scanning confocal microscope or an Olympus VS120 slide scanner. Images were captured with identical settings blinded to treatment groups and analysed using the trainable Weka segmentation plugin (Arganda-Carreras et al., 2017). A subset of images representative of the range of brightness/contrast of each individual immunohistochemistry image dataset were utilized for training and model generation, which was then applied to each dataset, respectively. Representative regions of interest were drawn around grey matter of individual images, then segmented binary images were analysed for count, percentage area of labelling, and density. These data were then used for statistical analysis.

### Statistical analysis

For all statistical analysis, generalised linear modelling and mixed effects models were used within the R statistical environment, as previously described (Collins et al., 2022). Initial analysis of mSOD1^G93A^ and C57BL/6 control groups was established to confirm that there was an alteration in measured factors based on disease. Drug treatment groups and animal sex were encoded as factors; ACY-738 positive/negative, and riluzole positive/negative, with a random intercept for each mouse for all statistical analysis. Generalised linear models were generated for continuous factors with motor function testing using the LME4 package (Bates et al., 2015). Non-continuous factors such as lower motor neuron counts, peripheral nerve size, and glial cell percentage area were analysed with the glmmTMB package (Brooks et al., 2017). Model performance was checked with the performance R package. Interactions for sex, age, ACY-738 treatment and riluzole treatment were performed for all behavioural analysis, while sex, ACY-738 and riluzole treatment were encoded as interactions and included for all histological analyses. Gaussian distribution modelling was performed for count-related data as previously described (Collins et al., 2022). Plots were generated using Prism v9.5 and ggpplot2 packages with either raw data + 95% confidence intervals, or conditional means ± 95% confidence intervals and beeswarm dots for individual animal values.

## Acknowledgements

The authors would like to thank Dr Thibaut Burg and Prof. Ludo Van Den Bosch for insight into drug treatment paradigms and collaboration, and to Dr Wesley J Olivier for the formulation of riluzole and ACY-738 at the University of Tasmania Chemistry Department.

## Authors’ contributions

AJP and SD both contributed equally to this work. SP and A.K contributed equally to this work.

---

**Supplementary Table 1:** Serum NFL levels are unaltered in mSOD1^G93A^ mice treated with ACY-738.

| *Predictors* | *Sex* | *N* | *Age (weeks)* | *Mean log*<br>*NFL(pg/uL)* | *95%CI* |
| --- | --- | --- | --- | --- | --- |
| mSOD1<sup>G93A</sup> vehicle | F | 8 | 14 | 6.81 | 6.50 – 7.13 |
| mSOD1<sup>G93A</sup> vehicle | M | 9 | 14 | 7.11 | 6.80 – 7.42 |
| mSOD1<sup>G93A</sup> riluzole | F | 8 | 14 | 6.77 | 6.46 – 7.09 |
| mSOD1<sup>G93A</sup> riluzole | M | 8 | 14 | 7.04 | 6.75 – 7.34 |
| mSOD1<sup>G93A</sup> ACY-738 | F | 9 | 14 | 6.52 | 6.19 – 6.85 |
| mSOD1<sup>G93A</sup> ACY-738 | M | 8 | 14 | 7.01 | 6.68 – 7.35 |
| mSOD1<sup>G93A</sup> ACY-738 + riluzole | F | 10 | 14 | 6.76 | 6.46 – 7.07 |
| mSOD1<sup>G93A</sup> ACY-738 + riluzole | M | 8 | 14 | 6.91 | 6.61 – 7.21 |
| mSOD1<sup>G93A</sup> vehicle | F | 8 | 20 | 8.41 | 8.12 – 8.70 |
| mSOD1<sup>G93A</sup> vehicle | M | 9 | 20 | 8.17 | 7.88 – 8.47 |
| mSOD1<sup>G93A</sup> riluzole | F | 8 | 20 | 8.04 | 7.74 – 8.34 |
| mSOD1<sup>G93A</sup> riluzole | M | 8 | 20 | 8.36 | 8.06 – 8.66 |
| mSOD1<sup>G93A</sup> ACY-738 | F | 9 | 20 | 7.89 | 7.61 – 8.18 |
| mSOD1<sup>G93A</sup> ACY-738 | M | 8 | 20 | 8.16 | 7.87 – 8.46 |
| mSOD1<sup>G93A</sup> ACY-738 + riluzole | F | 10 | 20 | 8.17 | 7.91 – 8.45 |
| mSOD1<sup>G93A</sup> ACY-738 + riluzole | M | 8 | 20 | 8.01 | 7.71 – 8.31 |

*Data presented as conditional log(mean) and 95% confidence intervals.*

